# Extinction recall of fear memories formed before stress is not affected despite amygdalar hyperactivity

**DOI:** 10.1101/261578

**Authors:** Mohammed Mostafizur Rahman, Ashutosh Shukla, Sumantra Chattarji

## Abstract

Stress is known to exert its detrimental effects not only by enhancing fear, but also by impairing its extinction. However, in earlier studies stress exposure invariably preceded both processes. Thus, compared to unstressed animals, stressed animals had to extinguish fear memories from higher levels of freezing caused by prior exposure to stress. Here we decouple the two processes to examine if stress specifically impairs fear extinction. Strikingly, when fear memories were formed before stress exposure, thereby allowing animals to initiate extinction from comparable levels of fear, recall of fear extinction was unaffected. Despite this we observed a persistent increase in theta activity in the BLA. Theta activity in the mPFC, by contrast, was normal. Stress also disrupted mPFC-BLA theta-frequency synchrony and directional coupling. Thus, in the absence of the fear-enhancing effects of stress, the expression of fear reflects normal regulation of mPFC activity, not stress-induced hyperactivity in the amygdala.

## Introduction

Accumulating evidence from animal models shows that stress elicits divergent patterns of plasticity across brain regions(Chattarji et al., 2015). For instance, repeated stress causes loss of dendrites and spines in the medial prefrontal cortex (mPFC)(Shansky and Morrison, 2009). In the basolateral amygdala (BLA), by contrast, chronic stress strengthens the structural basis of synaptic connectivity through dendritic growth and spinogenesis(Chattarji et al., 2015). Physiological and molecular measures of synaptic plasticity also exhibit these contrasting features. As useful as these studies have been in examining the effects of stress across biological scales, much of this evidence was gathered from postmortem analyses(Chattarji et al., 2015). Less is known about how stress affects neural activity in the intact brain of behaving animals. Further, in many of these studies, stress-induced plasticity was viewed as stand-alone effects intrinsic to individual brain areas, despite extensive interconnections between them. Indeed, interactions between these brain areas together give rise to behaviors that are affected by stress(Quirk and Mueller, 2008).

One such behavior involves the expression of fear memories, various facets of which depend on both the BLA and mPFC(Sierra-Mercado et al., 2011). Repeated stress has been shown to enhance fear memories, as well as impair their extinction(Miracle et al., 2006; Suvrathan et al., 2014). These studies, however, first exposed animals to stress, and then subjected them to fear conditioning followed by extinction(Izquierdo et al., 2006; Miracle et al., 2006). Consequently, stressed animals had to extinguish fear memories that were invariably stronger than unstressed animals, raising the possibility that the higher levels of freezing in stressed animals at the beginning of extinction contributed to the subsequent deficit in fear extinction (Zhang and Rosenkranz, 2013). One way to overcome this confound is for animals to form fear memories *before* stress exposure so that they can undergo extinction of fear from the same levels of freezing as their unstressed counterparts. This in turn would offer an opportunity to examine how stress specifically affects expression of fear after extinction, without any confounds of enhanced fear caused by prior exposure to stress. Hence, the present study combines simultaneous behavioral and *in vivo* electrophysiological analyses to address this question.

## Results

Rats were first subjected to auditory fear conditioning (Day 1, **Fig. 1**) at the end of which they exhibited significantly higher freezing behavior in response to the tone conditioned stimulus (CS) (**Fig. 1b**). These animals were then divided into two groups – one was subjected to 10 days of chronic immobilization stress (Days 2-11, **Fig. 1a**) while the other served as unstressed control. 24 hours after the end of chronic stress there was no difference in CS-induced freezing behavior between the two groups (Day 12, **Fig. 1b**). Thus, the recall of fear memory formed earlier was not affected by subsequent stress. This ensured that both stressed and unstressed animals were at the same levels of freezing when the extinction protocol was initiated after stress. Next, repeated tone presentations reduced freezing levels significantly such that both groups eventually underwent comparable extinction of fear, though the stressed rats were slower in achieving the same reduction in freezing (Day 12, **Fig. 1b**). Notably, a day later stressed animals showed no difference in recall of extinction memory compared to unstressed animals (Day 13, **Fig. 1b**). Thus, freezing levels during both fear and extinction recall were unaffected in stressed animals. This result differs from past reports of stress-induced deficits in fear extinction. However, as mentioned earlier, those earlier studies subjected animals to fear conditioning and extinction *after* exposure to stress (Maren and Holmes, 2016). Taken together, this suggests that the *timing* of stress may be a critical determinant of whether extinction recall is impaired. Thus, to examine this possibility we repeated the experimental design adopted in earlier studies by administering the same 10-day chronic immobilization stress protocol *prior* (**Fig. 1 Supplementary 1**) to the same fear conditioning paradigm depicted in **Fig. 1a**. Consistent with earlier findings, prior exposure to chronic stress caused a significant impairment in the recall of fear extinction (**Fig. 1 Supplementary 1a and c**). However, these animals, unlike those used in **Fig. 1** (i.e. conditioning before stress) were not implanted with electrodes for simultaneous *in vivo* recordings. Hence, we repeated the behavioral experiments described in **Fig. 1** without surgical interventions related to *in vivo* recordings. This too yielded the same results as seen in the implanted animals – extinction recall was intact in stressed animals (**Fig. 1 Supplementary 1b and d**) when they were subjected to conditioning before chronic stress. Thus, taken together these behavioral results suggest that stress-induced impairment in extinction recall is evident when stress is administered before, but not, after conditioning.

**Figure 1.**
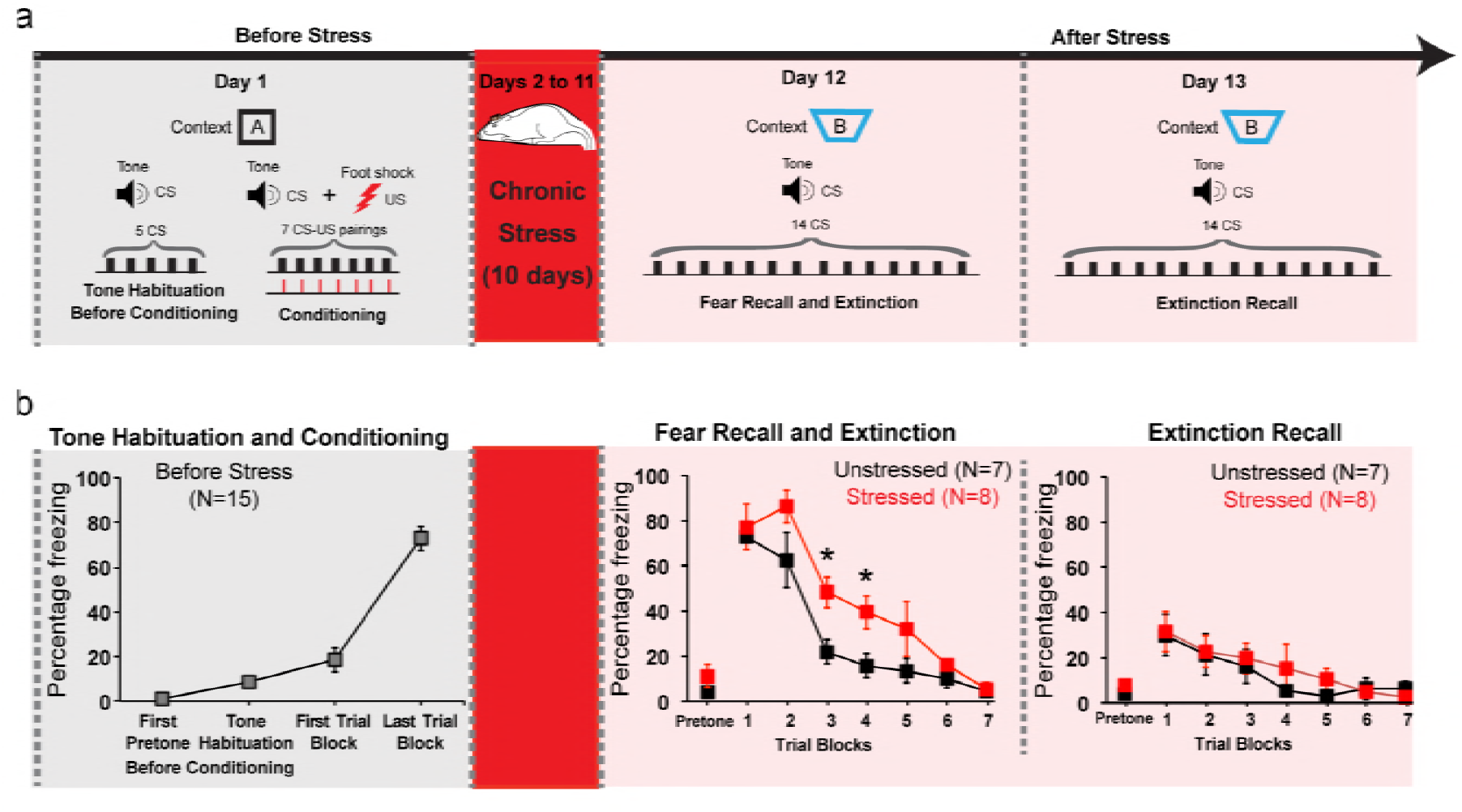
Normal expression of fear memories formed before stress. **(a)** Experimental design. Rats were subjected to tone habituation followed by fear conditioning on Day1. 24 hours later some of these rats were subjected to chronic immobilization stress (2h/d, 10d) while others were controls. Both groups underwent fear extinction 10 days later (Day 12), and extinction recall on Day 13. **(b)** Freezing at different time points. Significant increase in freezing relative to tone habituation after fear conditioning (*F*_(2.100,29.41)_=91.71, *P*<0.01). No difference in fear recall between stressed (N=8) and unstressed (N=7) rats (Day 12). Stressed rats exhibited higher CS-induced freezing in the third and fourth trial blocks (*P*<0.05) indicating delay in acquisition of fear extinction. However, both groups eventually decreased freezing to the same level at the end of the 7 trial blocks. There was no difference in freezing during extinction recall (*F*_(1,13)_=0.16, *P*=0.70). Data are mean ± s.e.m. in blocks of two trials except pretone. ^*^*P*<0.05.

Next, we examined the neural basis of this result by recording local field potentials (LFPs) in these freely behaving rats (Karalis et al., 2016; Likhtik et al., 2014). While a role for potentiation of amygdalar neuronal responses to the tone CS in conditioned fear behavior is well established, *in vivo* recordings have also shown correlations of tone responses in the dorsal mPFC (dmPFC) with freezing behavior in fear conditioning and extinction. Taken together with earlier pharmacological inactivation studies, these findings identified an important role for the dmPFC in underlying conditioned fear responses and the expression of fear extinction (Burgos-Robles et al., 2009; Sierra-Mercado et al., 2011). Therefore, in addition to the BLA, we also monitored responses in the dmPFC (**Fig. 2a, Supplementary 1**). We first analyzed CS-evoked LFPs in the BLA at three key behavioral time points described in Fig. 1 – tone habituation before conditioning, fear recall and extinction recall (**Fig. 2b-d**). During fear recall, auditory evoked potential (AEP) amplitudes(Rogan et al., 1997) were enhanced in both stressed and unstressed animals (**Fig. 2c, e**). However, while this increase was reversed in unstressed animals, it persisted in stressed rats even during extinction recall (**Fig. 2c, e**). Previous work also identified increase in CS-evoked theta power as a neural correlate of conditioned fear(Likhtik et al., 2014). BLA theta power in unstressed animals also paralleled the increase, followed by decrease, in freezing during fear and extinction recall respectively (**Fig. 2d-e**). By contrast, BLA theta power remained high in stressed animals (**Fig. 2d-e**). Thus, despite stress-induced hyperactivity in the BLA, fear expression was not enhanced during extinction recall. To probe this further, we also analyzed the same LFP parameters in the dmPFC, which according to recent studies plays an important role in fear expression(Karalis et al., 2016; Likhtik et al., 2014). In the dmPFC of unstressed rats, AEP amplitude increased during fear recall, and this was reversed during extinction recall (**Fig. 2e**). Stressed animals, however, did not exhibit any change in dmPFC AEP amplitudes during either fear or extinction recall. Further, fear conditioning enhanced dmPFC theta power in both stress and unstressed animals (**Fig. 2e**). Interestingly, this was reversed in both groups during extinction recall. In other words, unlike the BLA, changes in theta power in the dmPFC, during fear and extinction recall, were not affected by stress. Moreover, in stressed animals, bidirectional modulation of theta power in the dmPFC, but not the BLA, accurately mirrored the changes in freezing, a behavioral expression of fear.

**Figure 2.**
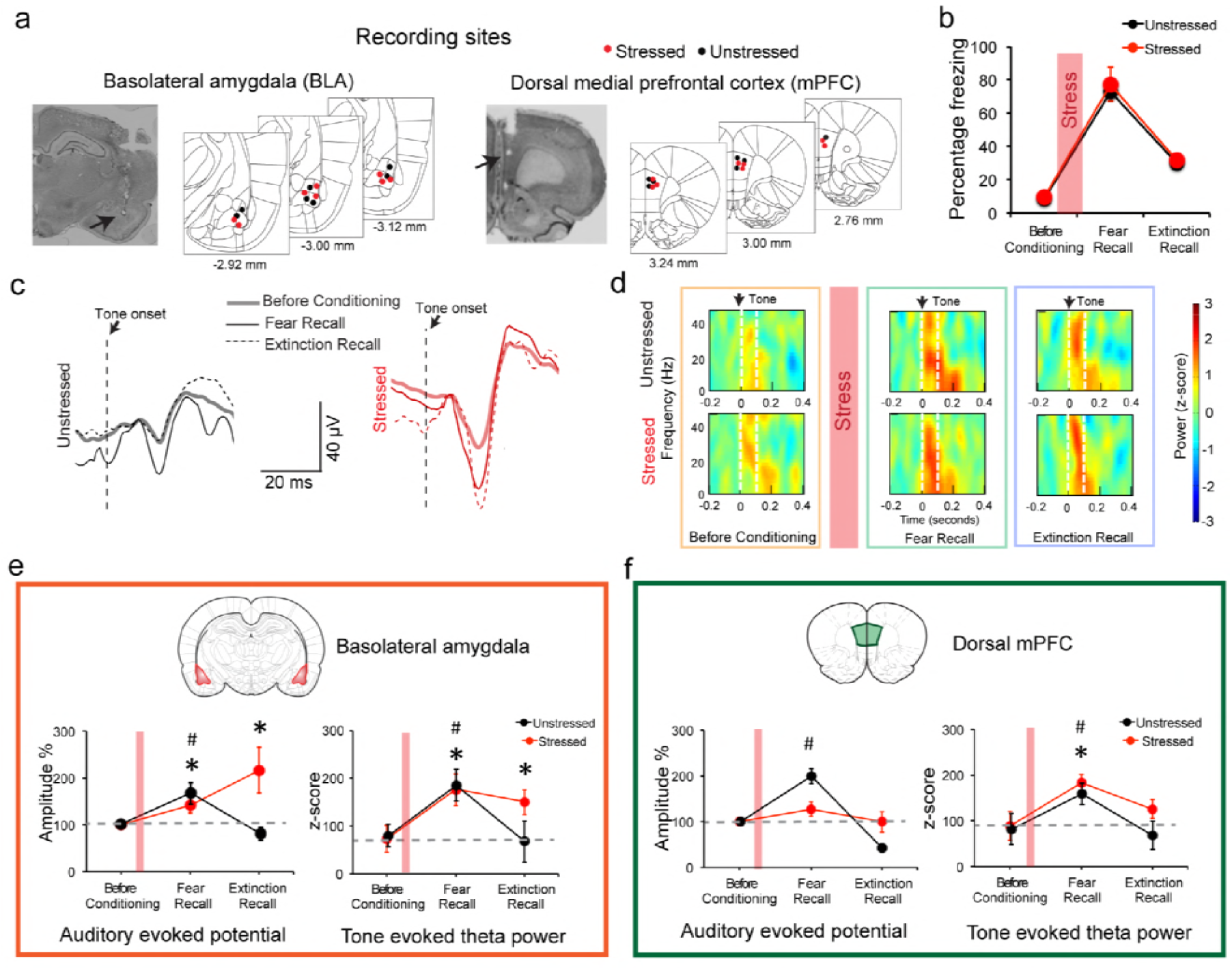
Stress triggered persistent hyperactivity in the BLA but not the mPFC. **(a)** Representative micrographs and diagrams of recording placements (red: stressed, black: unstressed) in the BLA (*left*) and mPFC (*right*). **(b)** Summary of changes in tone-induced freezing behavior before conditioning, and during fear and extinction recall. Stress (red bar) did not affect fear retrieval or extinction (Factor Stress: *F*_(1,13)=_0.26, *P*=0.62). **(c)** Representative BLA raw LFP traces depicting changes in AEPs recorded in response to CS presented during habituation before conditioning, fear and extinction recall in stressed (red) and unstressed (black) animals. **(d)** Representative spectrograms of BLA LFPs before conditioning (*left*), fear recall (*center*) and extinction recall (*right*) recorded from unstressed (top) and stressed (bottom) animals. Dotted white lines on the spectrogram indicate onset (arrow) and end of CS. **(e)** Percentage changes (normalized to tone habituation before conditioning) in auditory evoked potential amplitude (AEP, *left*) and auditory evoked theta power (*right*) in the BLA. All statistical comparisons are done within groups across the three time points, not between stressed (N=8) and unstressed (N=7) rats. In unstressed animals, AEP amplitudes increased during fear recall compared to tone habituation (*F*_(1.260,7.599)_=7.39, *P*=0.02) and was subsequently reversed during extinction recall. In stressed animals, by contrast, BLA AEP amplitudes were enhanced during both fear recall and extinction recall (*F*_(1.160,8.120)_=5.779, *P* =0.04). The same pattern of changes were observed in BLA theta power during fear recall in unstressed animals (*F*_(1.470,8.820)_=5.657, *P* =0.03), while stressed animals exhibited theta power enhancement during fear recall that persisted even during extinction recall (*F*_(1.957,13.70)_=13.70, *P*<0.01). **(f)** Percentage changes (normalized to tone habituation before conditioning) in AEPs (*left*) and auditory evoked theta power (*right*) in the dmPFC. All statistical comparisons are done within groups across the three time points, not between stressed (N=8) and unstressed (N=7) rats. AEP amplitudes in unstressed animals increased only during fear recall (*F*_(1.461,8.767)_=62.01, *P*<0.01) whereas the stressed animals exhibited no changes (*F*_(1.473,10.31)_=1.59, *P* =0.25). But dmPFC theta power increased in both unstressed and stressed animals during fear recall relative to tone habituation (Unstressed *F*_(1.223,7.336)_=5.894, *P*=0.04; Stressed *F*_(1.849,12.94)_=11.15, *P*<0.01). And these increases were reversed during extinction recall in both groups. Data are mean ± s.e.m. in each block; #*P***<**0.05, unstressed animals; \**P***<**0.05, stressed animals.

Finally, there is growing appreciation of the importance of interactions between the mPFC and BLA, not just activity within these areas, in regulating fear behavior(Karalis et al., 2016; Lesting et al., 2011; Likhtik et al., 2014; Popa et al., 2010). This issue comes into sharp focus here because of the distinct effects of stress on the mPFC versus BLA. Thus, in light of recent reports that theta frequency oscillations synchronize dmPFC–BLA circuits during expression of fear behavior(Karalis et al., 2016; Likhtik et al., 2014), we investigated whether the tone-evoked increases in theta power (**Fig. 2**) were accompanied by enhanced theta-frequency synchrony between the two areas, and if this was in anyway affected by stress. Hence, we quantified CS-evoked coherence to asses moment-by-moment synchrony across LFPs recorded from the dmPFC and BLA for all three time points (**Fig. 3a**)(Likhtik et al., 2014). In unstressed rats, consistent with earlier reports, the CS elicited significantly higher theta coherence during fear recall(Likhtik et al., 2014), and this increase persisted during extinction recall as well (**Fig. 3b**). Notably, in stressed animals, there was no change in BLA-dmPFC theta-frequency coherence (**Fig. 3b**). Thus, stress appears to completely suppress the dynamic, behaviorally relevant enhancement in BLA-dmPFC coherence that is seen during both fear and extinction recall in unstressed animals. In light of strong reciprocal connections between the mPFC and BLA, increases in theta synchrony have led earlier studies to analyze the direction of information flow between the two areas(Karalis et al., 2016; Likhtik et al., 2014). Hence, we estimated the directionality of functional connectivity and leads between the dmPFC and BLA using a previously validated method of calculating cross-correlations of instantaneous amplitude of filtered LFPs(Adhikari et al., 2010). This reveals that theta activity in the dmPFC leads that in the BLA during recall of both fear and extinction memories in unstressed rats (**Fig. 3c**). However, this dmPFC-to-BLA directional influence is absent in stressed animals. Together, these data suggest that chronic stress causes a decoupling of activity between the two brain areas, as evidenced by a complete disruption of the increase in dmPFC-BLA theta synchrony and dmPFC-to-BLA directional influence normally seen during the recall of fear and extinction memories.

**Figure 3.**
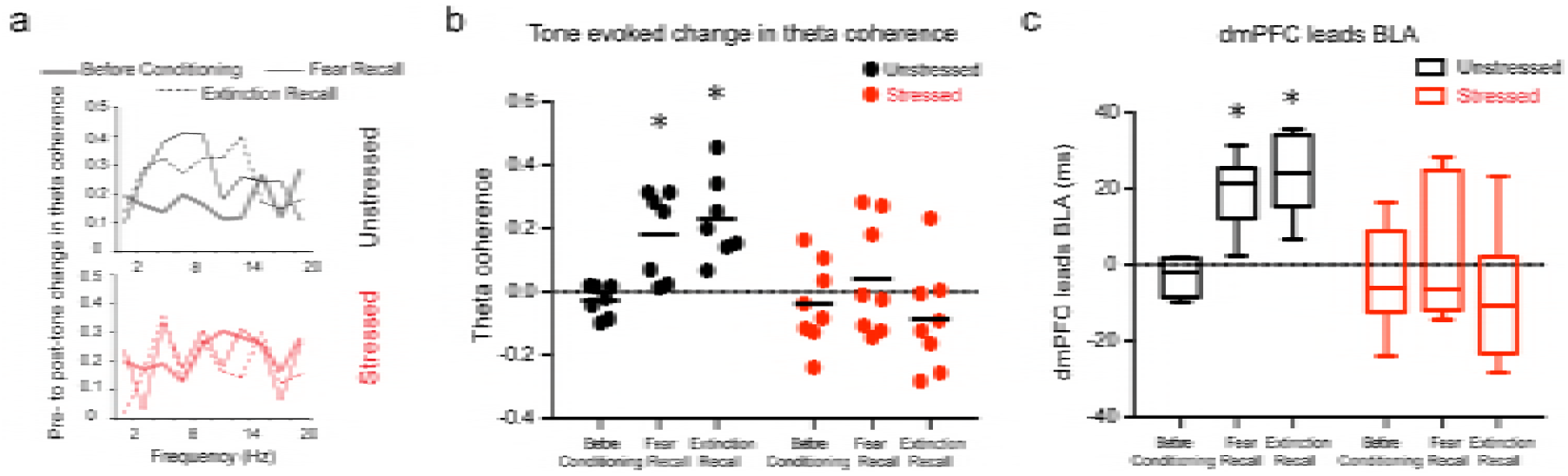
Stress disrupts enhanced mPFC-BLA theta synchrony and directional coupling during fear expression. **(a)** Tone-evoked changes in theta-frequency coherence between the BLA and dmPFC in exemplars of unstressed (*top*) and stressed (*bottom*) animals. Unstressed animals exhibited higher synchrony during fear recall and extinction recall than tone habituation (*F*_(1.408,8.451)_=10.66, *P*<0.01). These changes were absent in stressed rats (*F*_(1.582,11.07)_=1.01, *P*=0.38). **(b)** Means and distribution of tone-evoked changes in theta-frequency coherence for both groups (Unstressed N=7, Stressed N=8). \**P***<**0.05. **(c)** Estimation of leads between the dmPFC and BLA using the amplitude cross-correlation. The dmPFC leads over the BLA during fear and extinction recall in unstressed (N=7), but not stressed (N=8) rats. Data are presented as medians ± maxima/minima. \**P***<**0.05 significantly different from chance for each time point in each group.

This study, specifically designed to administer chronic stress after the formation of fear memory, reveals that when stressed animals started extinguishing fear memories from the same level of freezing as their unstressed counterparts, their ability to recall extinction memory remained intact. This is in contrast to past findings wherein stressed animals exhibited a deficit in extinction recall when faced with the challenge of extinguishing higher levels of fear caused by prior exposure to stress. Together, this suggests that the earlier findings were not the result of a deficit in extinction per se, but the higher initial levels of freezing before extinction. Interestingly, despite no visible behavioral effect of stress on fear expression, our *in vivo* recordings reveal a robust impact of stress on amygdalar activity, as evidenced by enhanced BLA theta activity that failed to reverse even after the animals exhibited normal extinction recall. This is consistent with earlier findings on physiological and structural strengthening of excitatory synaptic connectivity, as well as reduced inhibitory tone, in the BLA after stress (Suvrathan et al., 2014). In striking contrast, stress did not disrupt normal bidirectional modulation of mPFC theta activity, which in turn was reflected in normal freezing behavior during recall of fear and extinction. This is consistent with growing evidence for a pivotal role played by the mPFC in fear expression(Dejean et al., 2016; Sierra-Mercado et al., 2011). For instance, recent work has demonstrated strong correlations between mPFC theta-frequency oscillations and conditioning-induced freezing behavior(Likhtik et al., 2014). Furthermore, we find normal mPFC theta activity to be decoupled from the hyperactive BLA, possibly reducing the latter’s influence on fear expression. This is similar to a report that even a single episode of stress can weaken functional connectivity between the two areas measured by resting state fMRI(Liang et al., 2014). Indeed, such stress-induced disruptions in prefrontal-to-amygdala connectivity is also known to affect social interaction and anxiety-related behaviors in rodents(Adhikari et al., 2015; Hultman et al., 2016). Finally, it is interesting to note that the same chronic stress paradigm was previously shown to strengthen functional connectivity from the amygdala to the hippocampus(Ghosh et al., 2013), which undergoes stress-induced deficits similar to the mPFC (Arnsten, 2015; Chattarji et al., 2015; McEwen and Morrison, 2013). In other words, although both the hippocampus and mPFC undergo similar forms of stress-induced deficits, the impact of stress on their individual interactions with the amygdala are strikingly different. A better understanding of these divergent features of aberrant interactions distributed across the amygdala-mPFC-hippocampal network, not just those confined within each area, may offer new insights into therapeutic interventions against the cognitive and emotional symptoms of stress-related psychiatric disorder.

## Materials and methods

### Experimental Animals

Naïve 8-9 weeks old male Sprague-Dawley rats weighing 300–350 grams at the start of the experiment (National Centre for Biological Sciences, Bangalore, India) and housed in groups of two were used in the study. They were maintained on a 14-hour/10-hour light/dark cycle and had access to water and a standard diet *ad libitum*. All experiments were conducted in accordance with the guidelines of the CPCSEA, Government of India and approved by the Institutional Animal Ethics Committee of National Centre for Biological Sciences.

### Experimental design

The experimental design comprised of experimental procedures conducted over a 4 weeks period. The animals were handled for 2-3 days to familiarize with the experimenter. This was followed by a surgery to implant bundle of electrodes in the basolateral amygdala and the dorsal medial prefrontal cortex of the animals. The animals were allowed to recover for 7-10 days after surgery. Next, the animals were subjected to a 15-day behavioral paradigm with simultaneous recording of local field potentials (LFPs) during behavior (**Fig 1a**). The animals were initially habituated to the conditioning context on day −1 and day 0. Next on day 1 the animals were subjected to the tone habituation and fear conditioning protocol. Then on day 2 the animals were randomly allotted to the stressed or the unstressed groups. The animals in the stressed group were subjected to a 10-day chronic immobilization stress (CIS) from day 2 to day 11, where as the animals in the unstressed group were just handled once a day during the same period. Subsequently, on day 12, that is 24 hours after the end of CIS the animals were subjected to fear recall and extinction. On day 13 the animals were subjected to fear extinction recall session. After the end of the behavioral paradigm the animals were sacrificed and brains were collected for histological examination. To test fear memory formed after stress. For the behavior experiments without LFP recordings, the animals were handled for 2-3 days and then subjected to the behavior protocol. For fear memory formed prior to stress, the animals were subjected to tone habituation and conditioning on day 1 followed by 10 day chronic stress paradigm. Subsequently, the animals were subjected to fear recall and extinction on day 12 and extinction recall on day 13. For fear memory formed after stress, the animals were first subjected to a 10 day chronic stress paradigm followed by tone habituation and conditioning on day 11. Subsequently, the animals were subjected to fear recall and extinction on day 12 and extinction recall on day 13. The animals were randomly allocated to either stressed or unstressed groups.

### Surgical procedure

For recording LFPs from the dmPFC and BLA, rats were surgically implanted with formavar insulated nichrome wire (25 microns diameter) bundles unilaterally in the BLA and dmPFC (AM Systems, Carlsborg, WA, USA).

Rats were induced into anesthesia with 5% isoflurane (Forane, Asecia Queensborough, UK) and then maintained in anesthesia with 1.5-2% isoflurane. The level of anesthesia was regularly monitored throughout the procedure using the pedal withdrawal reflex to toe pinch. The animal was placed and head fixed on a stereotaxic frame. Body temperature of rats was maintained with a heating pad. Burr holes were drilled at the stereotactic coordinated of the BLA (stereotaxic coordinates were: 3.0 mm posterior to bregma and ±5.3 mm lateral to midline(Paxinos and Watson, 2009)) and the dmPFC (stereotaxic coordinates were: 3.0 mm anterior to bregma and ±0.5 mm lateral to midline(Paxinos and Watson, 2009)). A bundle of 8 formavar coated nichrome electrodes were then implanted using the stereotactic frame (8.3 mm and 3.4 mm ventral from the brain surface for BLA and dmPFC respectively). The implant was secured using anchor screws and dental acrylic cement. Rats were allowed to recover for 7–8 days following surgery. In the post-surgery period the animals were singly housed in separate cages. A total of 18 animals were implanted. Three animals were excluded from the study because the positioning of the electrode bundles was incorrect. The location of the cannulae placement for the 15 animals used in the study is shown in **Fig 2a.**

### Stress Protocol

Rats in the stressed group were subjected to a chronic immobilization stress (CIS) paradigm(Ghosh et al., 2013), consisting of complete immobilization for 2 hours per day (before noon) in rodent immobilization bags without access to either food or water, for 10 consecutive days.

### Fear Conditioning and Extinction protocol

Fear conditioning and extinction took place in different contexts placed inside sound-isolation boxes (Coulbourn Instruments, Whitehall, Pennsylvania, USA). Conditioning was performed in a box with metal grids on the floor (context A: 12 inches wide × 10 inches deep × 12 inches high, no odour). Fear extinction and extinction recall was performed in another context, a modified homecage (context B: 14 inches wide × 8 inches deep × 16 inches high, mint odour). Lighting conditions and walls were different between the two contexts. All chambers were cleaned with 70% alcohol before and after each experiment.

The behavior of the animals was recorded using a video camera mounted on the wall of the sound isolation box and a frame grabber (sampling at 30 Hz). The videos were analysed offline for further quantification of freezing behavior. Infrared LED cues were placed on the walls of the experimental chambers. These cues were activated in coincidence with auditory stimuli to monitor the tone-evoked freezing response offline. A programmable tone generator and shocker (Habitest system, Coulbourn Instruments, Whitehall, Pennsylvania, USA) were used to deliver tones and foot-shock during the experiment. Foot-shocks were delivered through the metal grids on the floor of the conditioning chamber. The tone was played using a speaker (4 Ω, Coulbourn Instruments, Whitehall, Pennsylvania, USA) placed inside the experimental chamber.

During context habituation, the animals were allowed to explore context A for 25 minutes in each session. Next, in the tone habituation session **(Fig 1a)** the animals received five presentations of an auditory tone (total duration of 30 seconds, 5 kHz auditory tone consisting of 30 pips of 100 millisecond duration at a frequency of 1 Hz; 5 millisecond rise and fall, 70 ± 5 dB sound pressure level) in context A. This was immediately followed by fear conditioning protocol, where the tone (CS) was paired (7 pairings, average inter-trial interval <ITI> = 120 seconds, with a range of 80–160 seconds) with a co-terminating 0.5 second scrambled foot shock (US; 0.7 mA). In the Fear Recall and extinction session the animals were presented with the same tone (CS) for 15 times (average inter-trial interval <ITI> = 120 seconds, with a range of 80–160 seconds) in the context B. Again, in the extinction recall session, the animals were subjected to the same CS 15 times again.

### Behavioral Analysis

Behavioral response was scored offline using video recordings of all the behavior sessions. Response to the auditory stimuli was evaluated in the form of freezing response. Freezing was defined as the absence of movement except due to respiration(Blanchard and Blanchard, 1988). The time spent freezing during the presentation of the tone was converted into a percentage score **(Fig 2c)**. The percentage freezing level was measured in every context/session for 30 seconds immediately before the presentation of the first tone trial to assess freezing in absence of an auditory stimulus. This was defined as the freezing in the pretone period **(Fig 2d)**. The pretone block represents freezing during the first pretone only. The tone habituation block represents freezing during the last two trials of tone habituation. The first and last trial blocks during conditioning represent the freezing during the first two and last trials of the fear conditioning session. The trial blocks in the fear recall and extinction session and the extinction recall session represent freezing over blocks of two trials each (1 to 14).

### In-vivo electrophysiological recordings

All the animals were subjected to the recording of the local field potentials (LFPs) during tone habituation session, fear recall and extinction session and extinction recall session. Auditory-evoked potentials (AEPs) were recorded by connecting the microelectrodes to a unit gain buffer head stage (HS-36-Flex; Neuralynx, Bozeman, Montana, USA) and a data acquisition system Digilynx (Neuralynx, Bozeman, Montana, USA). Neural data were amplified (1000 times) and acquired at a sampling rate of 1 kHz followed by a band-pass filter (1-500 Hz) using Cheetah data acquisition software (Neuralynx, Bozeman, Montana, USA).

### Data Analyses

#### Auditory Evoked Potentials (AEPs)

AEPs were averaged over all the tone pips for the specified trial blocks. Averaged AEPs were quantified by measuring the amplitude(Ghosh et al., 2013). The amplitude was measured by the difference between the maxima (dot) after the onset of the response and the negative peak (arrow) (**Fig 2 Supplementary 1**). AEP amplitudes were calculated before conditioning (last two trials of tone habituation), fear recall (first two trials of fear recall and extinction) and extinction recall (first two trials of extinction recall). The AEP amplitudes for all the sessions were normalized as a percentage to the AEP amplitude before conditioning for each animal.

### Time Frequency Analyses

Event related changes in spectral power were evaluated by time-frequency analysis performed using continuous wavelet transformation (MATLAB) on the averaged AEPs. Complex Morlet wavelets were used to compute phase and the amplitude of evoked responses within a frequency range from 2 to 100 Hz in steps of 0.1 Hz (Ghosh et al., 2013). The bandwidth parameter and center frequency of the mother wavelet were 2 and 1 Hz respectively. Subsequently, the wavelet power of the time series was calculated and expressed in decibels. This was followed by z-score calculation for each frequency band across the time series. Baseline average power for the duration of 0 to -200 milliseconds was subtracted across all time points for each frequency bands. Tone evoked theta power was calculated over the duration of 0 to 250 milliseconds from tone onset for frequency from 2 to 12 Hz.

### Coherence and Amplitude Correlation

Coherence between the BLA and dmPFC was calculated using Welch’s method. The window length used was 500 ms and 1,024 FFTs. Coherence was calculated over a wide frequency range. Theta coherence was quantified over 2-12 Hz frequency bandwidth.

For amplitude cross correlation, the signals were filtered between 2-12 Hz. Then instantaneous phase and amplitude were obtained using Hilbert transformation. Next the mean amplitude was subtracted from the instantaneous amplitudes to remove the DC component. Then the cross-correlation between the amplitudes of the two signals was computed with over lags ranging from +0.1 to **−**0.1 s. The lead/lag at which the cross-correlation peaked was then determined. Then the dmPFC and BLA instantaneous amplitudes were randomly shifted by 2-5 seconds relative to each other 100 times. The shifted amplitudes were cross-correlated to find peaks expected by chance. The actual cross-correlation was considered significant if its peak value was greater than 95% of the peaks generated by randomly shifted signal cross-correlations. Finally the distributions of these peaks were obtained and the time bin for the maxima of the distribution was quantified as the resultant lead/lag between the signals.

### Histology

After the experiment was concluded, rats were deeply anesthetized (ketamine/xylazine, 100/20 mg per kg). Electrolytic lesions (20 μA, 20 seconds) were made to mark the *in vivo* infusion sites. The animals were then perfused transcardially with ice-cold saline (0.9%) followed by 10% (vol/vol) formalin. The perfused brain was left in 10% (vol/vol) formalin overnight. Coronal sections (80 μm) were prepared using a vibratome (VT 1200S, Leica Microsystems, Wetzlar, Germany) and mounted on gelatin-coated glass slides. Sections were stained with 0.2% (wt/vol) cresyl violet solution and mounted with DPX (Sigma-Aldrich, St. Louis, Missouri, United States). The slides were imaged to identify and reconstruct infusion sites **(Fig 2a)**.

### Statistical Analyses

All values are expressed as mean + SEM unless mentioned otherwise. Each data set was evaluated for outliers, which was defined as greater than twice the standard deviation away from the mean. Freezing within the tone habituation and conditioning session was evaluated using a one way repeated measures ANOVA followed by Tukey’s post hoc test. Freezing in the fear recall and extinction as well as the extinction recall session was analysed using two ways repeated measures ANOVA followed by Holm-Sidak’s post hoc test. For the AEP amplitudes and tone evoked theta powers, one way repeated measures ANOVA followed by Tukey’s post hoc test was used to analyse the data for unstressed and stressed groups separately. Next, to determine if the theta coherence and amplitude correlation lead/lags are significantly different than zero, Student’s t test was used. For the behavior experiments without LFP recordings, two way repeated measures ANOVA was used to analyse the data for the tone habituation and conditioning session and the fear recall and extinction session. This was followed by Holm-Sidak’s post hoc test. For the extinction recall session unpaired Student’s t test was used. All the data sets passed the normality test. Sample sizes were not determined prior to the experiments and were sufficient to get significant effects. All statistical tests were performed using GraphPad Prism (GraphPad software Inc., La Jolla, California, USA).

## Acknowledgements

This work was supported by funds from the Department of Atomic Energy and Department of Biotechnology, Government of India, and the Madan & Usha Sethi Fellowship.

## Author Contributions

MMR and SC contributed to the experimental design. MMR and AS performed the experiments and analysed the data. MMR and SC interpreted the results. MMR and SC wrote the manuscript.

---

**Figure 1 Supplementary 1.**
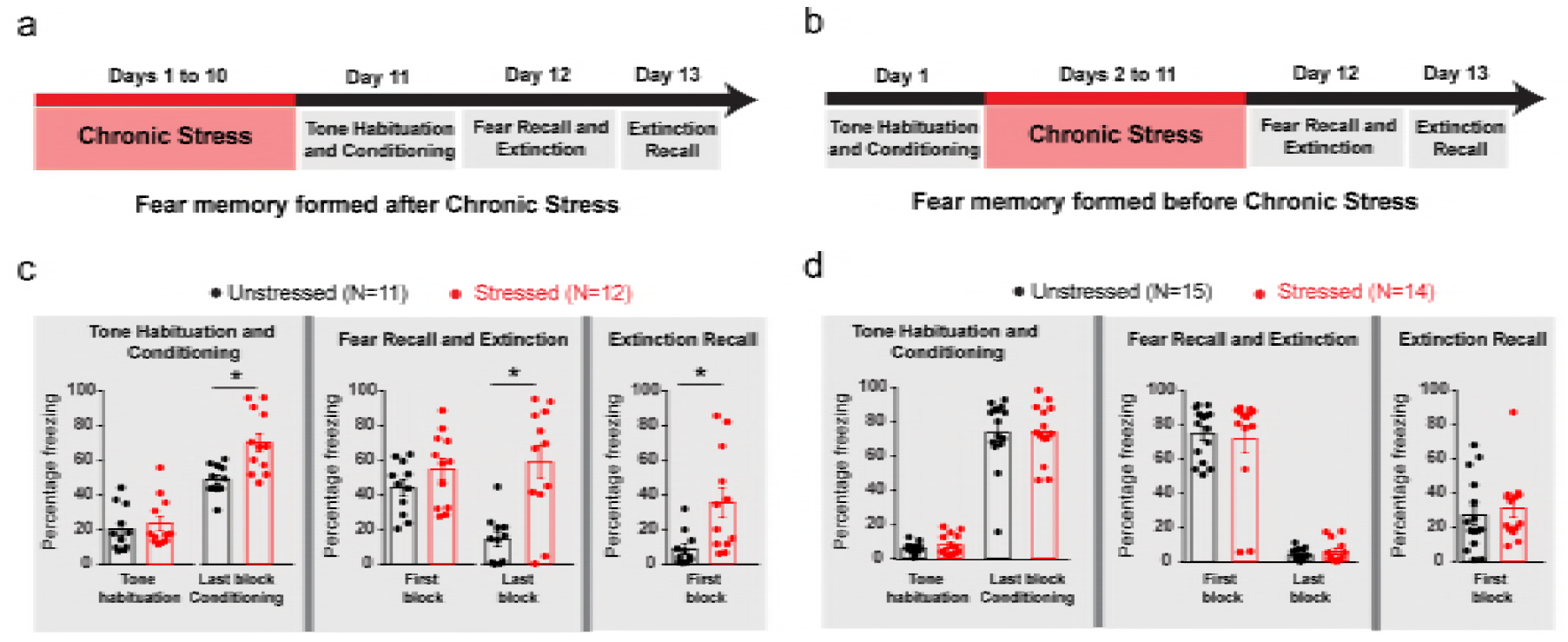
Impairment in the extinction of fear memories formed after, but not before, chronic stress. **(a)** Experimental design for testing the impact of chronic stress on fear memory formed *after* stress. Rats were subjected to chronic immobilization stress (2h/d, 10d) then subjected to tone habituation followed by fear conditioning on Day11. 24 hours later the animals underwent fear extinction (Day 12), and extinction recall on Day 13. A control group of unstressed rats underwent the same protocol but without chronic stress. **(b)** Experimental design for fear memory formed prior to the same 10-day chronic stress. Rats were subjected to tone habituation followed by fear conditioning on Day1. 24 hours later some of these rats were subjected to chronic immobilization stress (2h/d, 10d) while others were controls. Both groups underwent fear extinction 10 days later (Day 12), and extinction recall on Day 13. **(c)** Freezing at different time points. Significant increase in freezing relative to tone habituation after fear conditioning in both stressed (N=12) and unstressed (N=11) rats (Factor Conditioning: *F*_(1,21)_=108.20, *P*<0.01; Factor Stress: *F*_(1,21)_=7.04, *P*=0.01; Factor Interaction: *F*_(1,21)_=6.36, *P*=0.02). However, the stressed rats show enhanced freezing as compared to unstressed group after conditioning (*P*<0.01). Interestingly, the stressed and unstressed groups of rats show similar freezing response at the start of fear recall and extinction on Day 12. But at the end of the extinction session, only the unstressed animals show a decreased fear response (Factor Extinction: *F*_(1,21)_=3.90, *P*=0.06; Factor Stress: *F*_(1,21)_=16.56, *P*<0.01; Factor Interaction: *F*_(1,21)_=6.79, *P*=0.02). The stressed group also shows higher freezing response than the unstressed group at the start of extinction recall session on Day 13 (Unpaired t test, *P*=0.01). **(d)** All the animals showed an enhancement of freezing response due to fear conditioning prior to stress (Factor Conditioning: *F*_(1,27)_=431.20, *P*<0.01). No difference in freezing response during the fear recall and at the end of fear extinction between stressed (N=14) and unstressed (N=15) rats (Factor Extinction: *F*_(1,27)_=283.60, *P*<0.01; Factor Stress: *F*_(1,27)_=0.04, *P*=0.85; Factor Interaction: *F*_(1,27)_=0.42, *P*=0.52) (Day 12). Also, there was no difference in freezing during extinction recall on Day 13 (Unpaired t test, *P*=0.60). Data are mean ± s.e.m. in blocks of two trials. \**P*<0.05.

**Figure 2 Supplementary 1.**
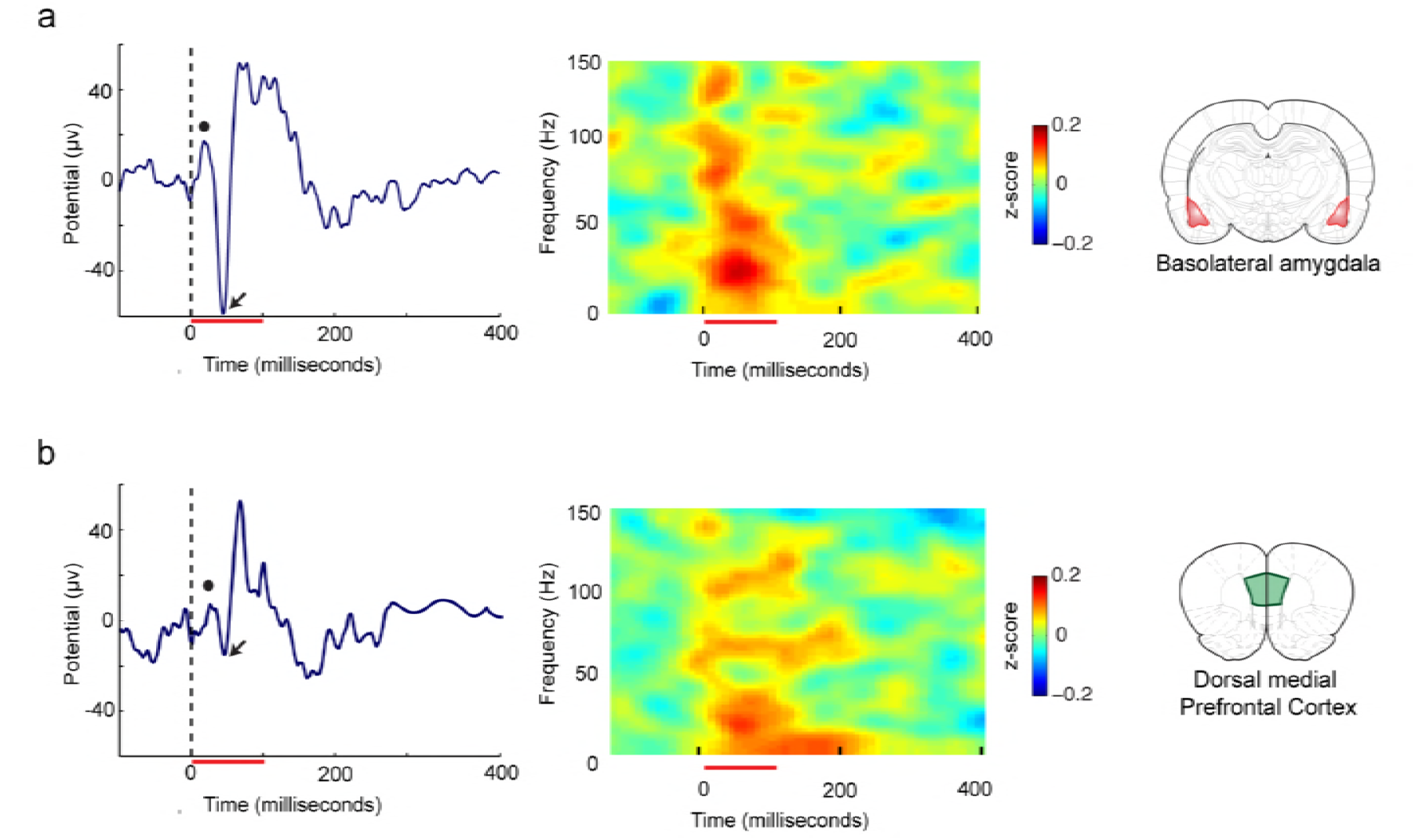
Example traces and power spectra of tone evoked local field potentials (LFPs). Tone induced average auditory evoked potential and tone evoked power spectra from the **(a)** basolateral amygdala (BLA), and **(b)** the dorsal medial prefrontal cortex. The red line at the bottom marks the duration of the CS tone. The black dot represents the first maxima and the arrow represents the first minima.

